# Genomic analysis for heavy metal resistance in *S. maltophilia*

**DOI:** 10.1101/404954

**Authors:** Wenbang Yu, Xiaoxiao Chen, Yilin Sheng, Qinghong Hong

## Abstract

*Stenotrophomonas maltophilia* is highly resistant to heavy metals, but the genetic knowledge of metal resistance in *S. maltophilia* is poorly understood. In this study, the genome of *S.maltophilia* Pho isolated from the contaminated soil near a metalwork factory was sequenced using PacBio RS II. Its genome is composed of a single chromosome with a GC content of 66.4% and 4434 protein-encoding genes. Comparative analysis revealed high syntney between *S.maltophilia* Pho and the model strain, *S.maltophilia* K279a. Then, the type and number of mechanisms for heavy metal uptake were analyzed firstly. Results revealed 7 unspecific ion transporter genes and 13 specific ion transporter genes, most of which were involved in iron transport. But the sulfate permeases belonging to the family of SulT/CysP that can uptake chromate and the high affinity ZnuABC/SitABCD were absent. Secondly, the putative genes controlling metal efflux were identified. Results showed that this bacterium encoded 5 CDFs, 1 copper exporting ATPase and 4 RND systems, including 2 CzcABC efflux pumps. Moreover, the putative metal transformation genes including arsenate and mercury detoxification genes were also identified. This study may provide useful information on the metal resistance mechanisms of *S.maltophilia.*

## Introduction

Heavy metals are naturally occurring elements with high toxicity to all kinds of lives. However, with the broad application of heavy metals, they have become a major kind of pollutants in soils and water bodies. These pollutants in the environment will finally accumulate in humans through the food cycle and cause serious diseases. Thus, to remove the heavy metals in the environment is very important. In recent years, the potential of microbes in removal of heavy metals is receiving more and more attention. To date, various microorganisms such as bacteria (Stahl *et al*, 2015), yeast (Wu *et al*, 1993), fungi (Mehra and Winge, 2010), and algae (Philippis *et al*, 2011) have been reported to tolerate and remove heavy metals from aqueous solutions.

*Stenotrophomonas maltophilia*, formerly named *Pseudomonas maltophilia*, is an aerobic, non-fermentative gram negative bacterium ubiquitous in the environment (Palleroni *et al*, 1993). This bacterium is receiving more and more attention as it is an important opportunistic pathogen, which exhibits intrinsic resistance to various antibiotics and is difficult to cure (Pages *et al*, 2008). However, this strain also has a broad range of metabolic activities which is related to great potential of biotechnology uses. First of all, it has promising uses in farming because of its powerful capabilities of recycling nitrogen/sulfur, promoting plant growth and inhibiting the growth of plant pathogens (Ryan et al, 2009). Then, it can degrade a lot of natural and man-made pollutants, including p-nitrophenol, 4-chlorophenol, polycyclic aromatic hydrocarbons, selenium compounds, benzene, toluene, ethylbenzene and xenobiotics (Ryan et al, 2009). Importantly, it also exhibits intrinsic resistance to various heavy metals, making it potential to be used in removal of metal pollution (Ryan et al, 2009).

The studies of Pages *et al*. have reported that the Sm777 strain of *S. maltophilia* could tolerate high levels (0.1-50 mM) of various metals, including the most toxic metals, Cd and Hg (Pages *et al*, 2008). According to their study, this strain could detoxify Cd to CdS (Pages *et al*, 2008). Besides, they reported that this strain could reduce high concentration of selenite and tellurite to elemental state (Pages *et al*, 2008). The studies of Blake *et al.* have also reported that *S. maltophilia* O2, isolated from soil at a toxic waste site, could catalyze the transformation and precipitation of numerous high level toxic cations and oxyanions (Blake *et al*, 1993). These results indicated that *S. maltophilia* might had various mechanisms to detoxify heavy metals. To date, many strains of *S. maltophilia* have been isolated and sequenced genomically. However, the genetic knowledge of the metal resistance mechanisms in *S.maltophilia* is still poorly understood. Thus, to get more insight into the different mechanisms of heavy metal tolerance in *S.maltophilia* would be of great interest to perform a genome analysis of this species (Pages *et al*, 2008). In this study, a complete genome sequence of *S. maltophilia.* Pho was reported to reveal the heavy metal tolerance mechanisms in *S. maltophilia*. This report may improve our understanding of the genetic basis of metal resistance in *S. maltophilia.* To our knowledge, this is the first study on the metal resistance of *S. maltophilia* at the genome scale.

## Materials and methods

### Bacterial isolation and DNA Extraction

The Pho strain was isolated from the heavy-metal contaminated soil near a metalwork factory using phenol as the sole carbon source (K_2_HPO_4_ 0.4 g/L, NaCl 0.1 g/L, (NH_4_)_2_SO_4_ 0.04 g/L, MgSO_4_ 0.12 g/L, MnSO_4_ 0.012 g/L, Fe_2_(SO_4_)_3_ 0.012 g/L, NaMoO_4_ 0.012g/L, Phenol 1 g/L). The culture was inoculated to agar plate using streak plating at 37°C. The strain was purified thrice. Three 500 ml exponential phase culture was collected and centrifuged at 10000 rpm for 30 min. The pelleted cells were washed with ddH_2_O twice. Then, the total DNA was extracted using DNA Kit (JieRui, Shanghai) according to the manufacturer’s instructions.

### Library Construction and DNA Sequencing

The purified total DNA was detected using agarose gel electrophoresis and Nanodrop2500 (OD260/280=1.8-2.0,>10μg). Then the genomic DNA was fragmented to prepare 8-10 Kb PacBio SMRTbell libraries using G-tubes. Both the ends of a fragment were added a single circle DNA. Then the singe circle DNA was fixed to the polymerase of ZMW (zero-mode waveguides). The genome was sequenced using the PacBio RSII platform. Genome sequencing was conducted at the Shanghai Meiji Biotechnology Co., Ltd. The genome sequence data were deposited in GenBank with the accession number CP029759.

### Genome Assembly, Annotation and Comparative Analysis

Low quality reads were filtered to get high quality reads. A total of 163103 reads with an average length of 7224.9 bp were obtained. The post filter reads length and quality distribution is shown in the Figure S1. Genome assembly was conducted according to hierarchical genome-assembly process (HGAP v2.3.0) assembly method. rRNA and tRNA were identified using the method of Barrnap 0.4.2 and tRNAscan-SE v1.3.1, respectively. Then gene prediction, genome annotation and genome island identification were conducted using the method as described in the reference (Chen *et al*, 2018). The genome comparisons between *S. maltophilia.* Pho and *S. maltophilia* K279a were conducted using the same genome annotation method as described above (Chen *et al*, 2018). Genomic Synteny between Pho and K279a was analyzed using progressive Mauve with a minimum LCB weight of 100 (Darling *et al*, 2004; Darling *et al*, 2007). The genomes were also compared by determining the numbers of genes that are unique to one organism and the number that are shared. Protein identical sites were defined by BlastP hit with a cutoff value of 60% identity.

### Data availability

Strains are available upon request. The genome sequence data were deposited in GenBank with the accession number CP029759. Supplemental Material, Figure S1 and Figure S2 show PacBio read length and quality distribution for the sequenced *S. maltophilia* Pho genome, respectively. Table S1 shows prediction and annotation of the encoding-genes in the genome of *S. maltophilia* Pho. Table S2 shows the assemble results of the genome sequence. Table S3 shows COG functional categories of this strain. Table S4 shows the statistical results of the genomic islands predicted in this genome. Table S5 shows the comparison results of protein identity between *S. maltophilia* Pho and *S. maltophilia* K279a. Table S6 shows proteins involved in heavy metal import. Table S7 shows the proteins involved in resistance of divalent metals. Table S9 shows the proteins involved in detoxification of arsenate and mercury.

## Results

### General overview

Genome of *S. maltophilia* Pho was sequenced using Pac-Bio RSII and a total of 163103 reads with an average length of 7225 bp were produced. The complete genome is composed of a 4,555,564 bp single chromosome with 66.4% GC (Figure 1). In this genome, 4434 protein-encoding genes with an average length of 891 bp were predicted. 2972 of the 4434 genes were assigned to 24 COG categories (Figure 1 and Table S3). The non-coding region contains 73 tRNAs and 13 rRNAs (5 5S rRNA, 4 16S rRNA, 4 23S rRNA) (Txt S1 and Txt S2). 16S rRNA sequence analysis confirmed that this bacterium is *Stenotrophomonas maltophilia.* Genomic island analysis was performed and 56 islands were predicted (Table S4). Moreover, 58 tandem repeats with a total length of 17089 bp were identified. Genomic data were submitted to Genebank with the accession number CP029759.

**Figure 1.**
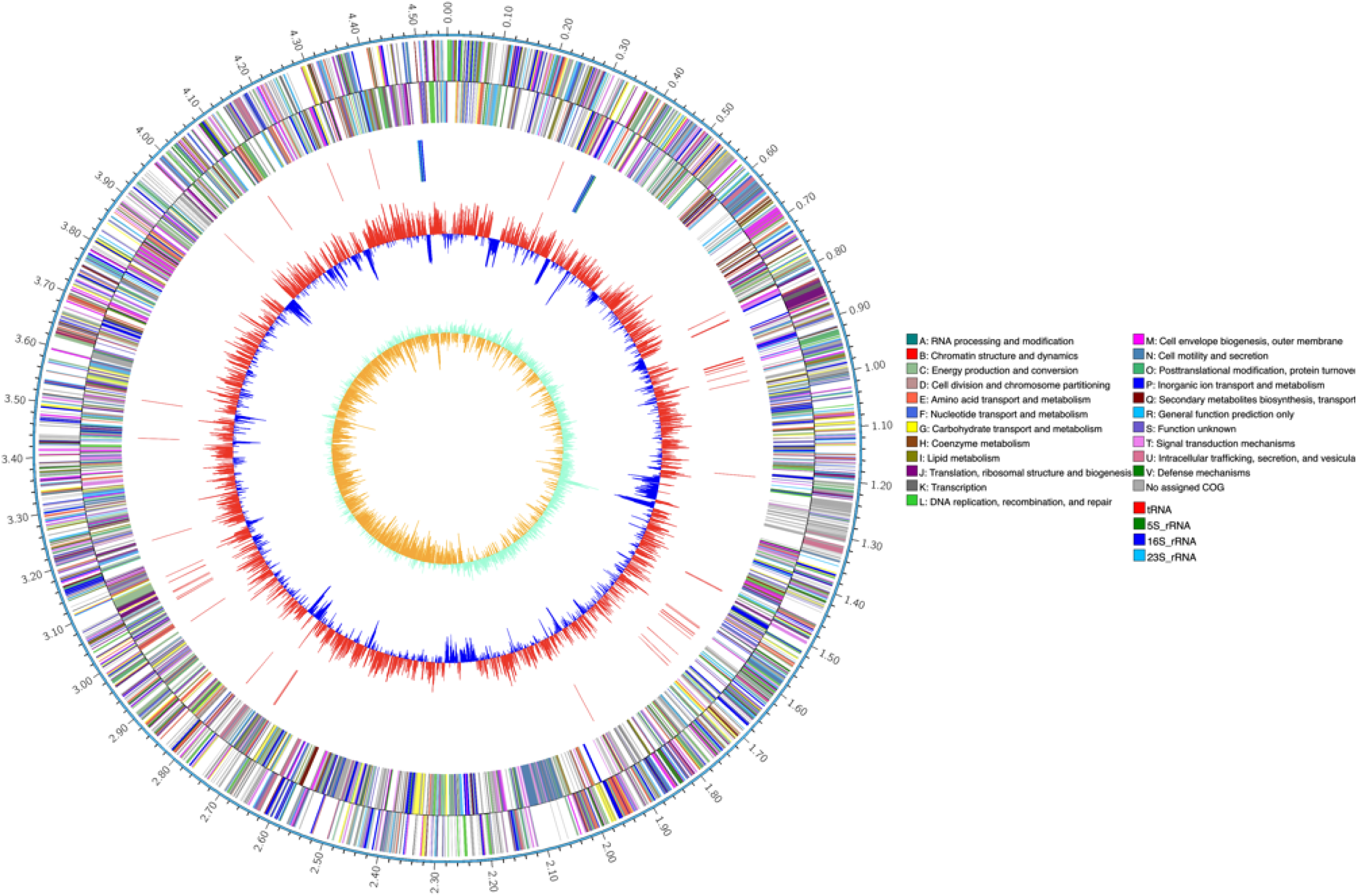
Chromosome atlas of *S.maltophilia Pho. dnaA* at position 1. From the outer to the inner circles. The first circle represents the size of the chromosome, with a degree scale of 0.1Mb. The second and third circles represent the CDS on the forward and reverse DNA chain, respectively. The colors on the second and third circles represent COG categories. The fourth circle represents the tRNA and rRNA. The GC contents are represented on the fifth circle, on which the red parts represent higher GC contents than the average level and the blue parts represents lower GC contents than the average level. The sixth circle represents the value of GC skew.

### Comparative analysis

The chromosome sequence of *S. maltophilia* Pho was compared to that of the model starin, *S. maltophilia* K279a. The chromosome of Pho was about 300 Kb shorter than K279a. Furthermore, there were 4434 and 4483 protein-encoding genes predicted in Pho’s and K279a’s genome using the same method, respectively. Progressive Mauve Alignment was used to compare the syntney between these two chromosomes and the results revealed extensive consensus identity (Figure 2). With a minimum LCB weight of 100, only 30 LCB blocks were identified in the comparison. Two translocation+inversions with a length of about 25.9Kb and 53.7Kb respectively were observed in the Pho’s genome (Figure 2). In the genome of Pho, about 4.2Mb were located in the LCB blocks, indicating about 4.2Mb nucleotides of the Pho genome has high homology to K279a and about 0.3Mb nucleotides of the Pho genome lack homology to K279a. In the Pho’s LCB blocks, a few complete white regions which represent specific elemental sequence were observed. Finally, all proteins of Pho were also compared to the proteins of K279a to determine the numbers of genes that are unique to Pho using BlastP method with a cutoff value of 60% identical sites. By that criterion, 575 proteins were found to be unique in Pho (Table S5).

**Figure 2.**
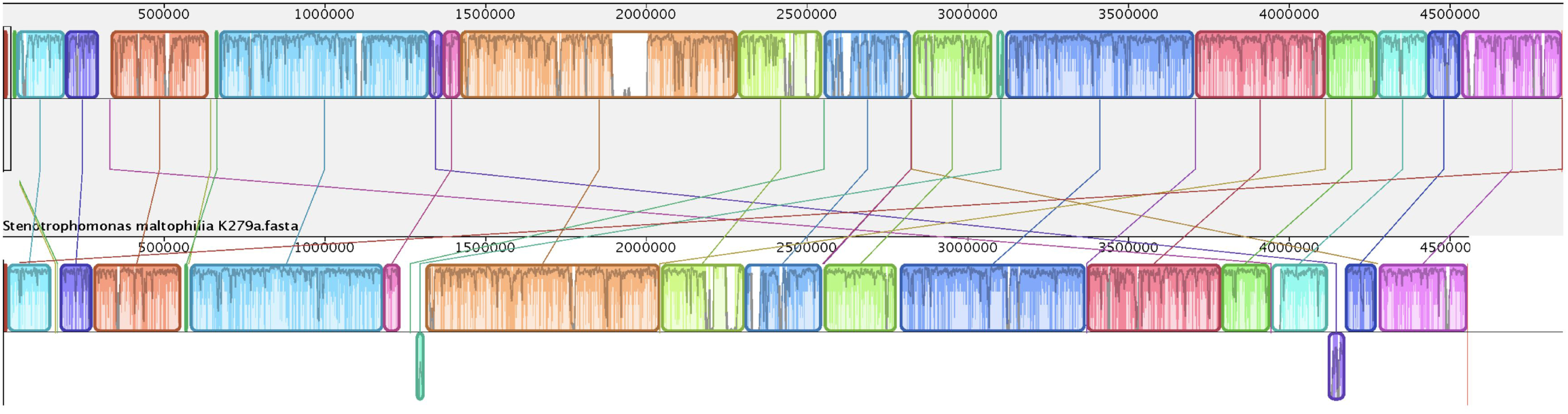
Collinearity analysis of the complete genome between *S.maltophilia* strain Pho and K279a. The upper and lower shafts represent the Pho and K279a genomes, respectively. Regions outside blocks lack detectable homology among the input genomes. Colored blocks in the first genome are connected by lines to similarly colored blocks in the second genome. These lines indicate which regions in each genome are homologous. Areas that are completely white were not aligned and probably contain sequence elements specific to a particular genome.

### Metal uptake systems

The first factor that determine the extent of metal resistance in bacteria is the type and number of metal uptake systems, through which heavy metal ions enter the cell. Bacteria often use two types of uptake system for heavy metal ions (Nies, 1999). The first is fast but unspecific and could accumulate nonessential toxic metal ions (Nies, 1999). 7 unspecific uptake systems were found in this genome. Sulfate permeases belonging to the family of SulT, SulP and CysP, could also uptake oxyanions such as chromate, molybdate, selenate and tungstate (Nies, 1999; Barajas *et al*, 2011; Nies and Silver, 1995). Herein, a protein of the SulP family, which also uptake selenate or selenite, was found (Table S6). However, proteins belonging to the family of SulT and CysP, were not found. The low affinity phosphate uptake protein, PitA, was responsible for arsenate accumulation (Nies, 1999). Herein, PitA was predicted to be encoded by STM1584 (Table S6). Low-affinity divalent cation transporters can uptake many divalent cations (Nies, 1999). The CorA is a Mg^2+^/Co^2+^ importer which may also accumulate Mn^2+^, Ni^2+^, Zn^2+^ and Cd^2+^; MntH is a Mn^2+^ transporter which may also accumulate Cd^2+^, Co^2+^, Zn^2+^ and to a lesser extent Cu^2+^ and Ni^2+^; while ZupT is a Zn^2+^ uptaker which may also accumulate Cu^2+^ and Cd^2+^ (Nies, 1999). Herein, 3 genes were predicted to encode CorA, MntH and ZupT respectively (Table S6). Moderate affinity Mg^2+^ transporter MgtA, MgtB and MgtE also uptake Mn^2+^, Zn^2+^ and Co^2+^ (Nies, 1999). Two genes were predicted to encode MgtA and MgtE, respectively (Table S2).

The second type of heavy metal uptake system is slower but has high specificity since it often uses energy of ATP hydrolysis (Nies, 1999). 13 genes were related to high-affinity heavy metal transporters (Table S6). 10 of them were annotated as high affinity Fe^3+^-siderophore receptor protein (Table S6). The other 3 were annotated as molybdate ABC transporter (Table S6). Surprisingly, high affinity uptake systems for Zn^2+^ (ZnuABC) and Mn^2+^ (SitABCD), which are encoded in a lot of gram-negative bacteria, were not found in this genome (Porcheron et al, 2013).

### Resistance of divalent heavy metals

In bacteria, resistance to divalent heavy metals, including Cd^2+^, Zn^2+^, Co^2+^, Mn^2+^ and Ni^2+^, are mainly dependent on the metal efflux systems, including cation diffusion facilitators (CDFs), P-type ATPases and resistance, nodulation, cell division (RND) efflux pumps (Nies, 1999; Jaroslawiecka and Piotrowskaseget, 2014). In the genome of Pho, 5 CDF transporters and 4 RND systems were predicted (Table S7). Of the 5 CDF transporters, STM0420 and STM1331 were annotated as CzcD (Table S7), which mediates resistance of low level Zn^2+^, Co^2+^, Cd^2+^ and Ni^2+^ (Wei Y and Fu D, 2005; Salusso A and Raimunda D, 2017). Moreover, there was a probable Mn^2+^ efflux pump, MntP, predicted in this genome (Table S7). STM0146, STM0147 and STM0148 were predicted to compose the CnrABC system, which mainly efflux Ni^2+^ and Co^2+^. In the up-stream region of STM0146, a Cnr regulator CnrR and RNA polymerase sigma factor CnrH, were predicted to be encoded by STM0144 and STM0145, respectively. All of these genes were located on the genomic island 2 with a length of 28.8 Kb (Table S7), indicating this gene cluster may be acquired from lateral gene transfer. CzcCBA, which efflux Zn^2+^, Co^2+^ and Cd^2+^, contains the inner membrane efflux pump CzcA, which is in complex with the membrane fusion protein CzcB and the outer membrane factor (OMF) CzcC (Stahl et al, 2015). Two genes (STM2188 and STM2420) were predicted to encode CzcA (Table S7). In the down-stream region of STM2188, STM2189 and STM2190 encode CzcB and CzcC respectively; while in the down-stream region of STM2420, two RND efflux pump periplasmic adaptor subunit and a CzcC protein were predicted. MdtABC is a recently identified new RND efflux pump of zinc (Porcheron et al, 2013). Herein, MdtABC was predicted to be encoded by STM3991-STM3993 (Table S7).

Metal sequestration is a mechanism to prevent exposure of essential cellular components to metals, commonly Cd^2+^, Zn^2+^ and Cu^2+^, by accumulating them within the cytoplasm with metallothionein or cysteine-rich proteins (Burins et al, 2000). However, there was no such metallothionein or cysteine-rich protein. Pages *et al*. have reported that *S. maltophilia* sequestrated Cd^2+^ by formation of Cd-S clusters (Pages et al, 2008). They suggested that Cd^2+^ might be sequestrated by sulfide produced from desulfhydrylation of cysteine (Pages et al, 2008). But, no homolog of cysteine desulfhydrase was found. Thus, the mechanism of Cd sequestration needs further characterization.

### Resistance of copper

Copper resistance involves of Cu^+^ efflux and oxidation in bacteria (Nies, 1999). There are two types of Cu^+^ efflux systems in gram-negative bacteria: one is the CusABCF RND efflux pump and the other is the copper exporting P-type ATPase. The CusABCF system was not found. However, two genes (STM1908 and STM2170) were annotated as copper P-type ATPase (Table S8). STM1908 was more identical to copper transporting ATPase ActP2, while STM2170 was more identical to copper exporting ATPase ActP (Data not shown). Thus, STM2170 may be involved in Cu^+^ efflux in the Pho strain. Multi-copper oxidase is important in detoxification of Cu^+^ by oxidizing it to Cu^2+^ (Porcheron et al, 2013). Herein, two multi-copper oxidases (MCOs) were found (STM2179 and STM3351, Table S8). In the down-stream region of STM2170, STM2172, STM2173, STM2175 and STM2178, were annonated as copper resistance protein PcoD, PcoC, CopG and PcoB, respectively. STM2174 was predicted to encode a copper storage protein, four-helix bundle copper-binding protein. STM2170-STM2179 were located on the island 27 with a length of 9.9Kb, indicating this cluster was probably acquired from lateral gene transfer.

### Resistance of Metalloid and Mercury

Microorganisms can detoxify heavy metal or metalloid species by metal transformation, including oxidation, reduction, methylation and dealkylation (Bruins, 2000). Herein, several genes were found to involve in metal reduction, but no gene was found to involve in metal oxidation, methylation or dealkylation (Table S9). Detoxification of arsenate involves of reduction of arsenate and efflux of arsenite [19]. STM0134 was predicted to encode arsenate reductase, ArsC (Table S9). STM0134 was placed in an arsenate detoxification gene cluster. In the upstream region of STM0134, STM0133 encoded arsenite efflux protein (Yun et al, 2000). In the downstream region of STM0134, STM0135 encoded an ArsR regulator. STM0136 encoded ArsH, which may be involved in high-level arsenate resistance (Branco et al, 2008).

Hg^2+^ detoxification involves importing of Hg^2+^ to the cytoplasm and reduction of Hg^2+^ to Hg, which will leave the cell (Nies, 1999). STM2143 was predicted to encode Hg^2+^ reductase and was placed in a mercury detoxification gene cluster (STM2140-STM2145) (Table S9). STM2140 encoded Hg^2+^ resistance operon regulator MerR. STM2142 and STM2141 encode MerP and MerT, which mediate transport of Hg^2+^. While STM2145 encoded a broad mercury transporter, MerE, which mediates transport of both Hg^2+^ and CH_3_Hg(I). STM2144 encodes a co-regulator of mercury operon regulator, MerD (Mukhopadhyay et al, 1991). However, MerB and MerC that are contained in mercury resistance operon of some bacteria were not present in the genome (Nies, 1999).

## Discussion

The first factor that determine the extent of metal resistance is the type and number of mechanisms for metal uptake (Burins, 2000). So we firstly identified the metal uptake systems. In this study, 7 unspecific uptake systems were found. These systems can mediate uptake of many toxic metal ions except chromate, since the CysP or SulT systems for chromate uptake were not present (Barajas et al, 2011). Thus, this strain of *S. maltophilia* will very likely not accumulate chromate in the cytoplasm. This may be supported by the absence of chromate efflux protein ChrA in the genomes of Pho and K279a (Data not shown). There were 13 genes found to be related to high-affinity heavy-metal uptake systems, but the ZnuABC and MntABC system were not present. ZnuABC is a high-affinity zinc uptake system which is required for growth in zinc limitation conditions and contributes to the virulence of pathogens (Ammendola et al, 2007); while MntABC is a high-affinity manganese uptake system which is required for growth in manganese limitation conditions and also contributes to the virulence of pathogens (Kehl-Fie et al, 2013). Since *S.maltophilia* can infect human bodies in which zinc and manganese is very limited, there may be alternative high-affinity zinc or manganese uptake mechanisms in *S.maltophilia*, as was seen in the strain of *Pseudomonas aeruginosa*, in which mutation of ZnuABC only result in slight growth defect even in zinc limited conditions (Porcheron et al, 2013).

Metal efflux systems are major mechanisms involved in resistance of heavy metals. Totally 11 efflux systems were identified, including 5 CDFs, 4 RNDs, a copper exporting ATPase and a manganese efflux protein. Most of these systems are involved in efflux of Zn^2+^, Co^2+^, Cd^2+^ and Ni^2+^. Probably, two CzcABC system were encoded in this genome. The first is encoded by STM2188-2190, which are located on the gene island 28 and 29. The second is probably encoded by STM2420-STM2423. STM2420 and STM2423 encode CzcA and CzcC respectively. STM2421 and STM2422 were predicted to encode RND efflux pump periplasmic adaptor subunits. Furthermore, these two proteins were also annotated as CzcB in the KEGG database (Table S1). Thus, we suggested that STM2420-STM2423 also composed CzcABC like system. In some strains of *S.maltophilia*, a Cd^2+^ P-type ATPase CadA was found, but this protein was not present. Pages *et al*. have reported that *S. maltophilia* may produce hydrogen sulfide from cysteine to sequestrate Cd^2+^ by formation of Cd-S clusters (Pages et al, 2008). Production of hydrogen sulfide from cysteine may be mediated by D-cysteine desulfhydrase, L-cysteine desulfidase and cystathionine lyase. Neither D-cysteine desulfhydrase nor L-cysteine desulfidase was present in this genome, but a cystathionine beta-lyase was found. Thus, cystathionine beta-lyase may be involved in Cd^2+^ sequestration in *S.maltophilia*.

*S.maltophilia* had a broad capability to transform numerous toxic metals and oxyanions. Arsenate and mercury reductase have been identified in this genome of Pho. The report of Blake *et al* has revealed that *S.maltophilia* could not reduce Se(V) and Te(V), but it could reduce selenite and tellurite (Blake et al, 1991). According to their study, selenite and tellurite were reduced by glutathione reductase in the cytoplasm. Herein, only STM3111 was predicted to encode glutathione reductase in the genome (Table S1).

